# Induction via Functional Protein Stabilization of Hepatic Cytochromes P450 upon gp78/Autocrine Motility Factor Receptor (AMFR) Ubiquitin E3-ligase Genetic Ablation in Mice: Therapeutic and Toxicological Relevance

**DOI:** 10.1101/623041

**Authors:** Doyoung Kwon, Sung-Mi Kim, Peyton Jacob, Yi Liu, Maria Almira Correia

**Author notes:** **Corresponding author:** M. A. Correia, Cellular & Molecular Pharmacology, Genentech Hall, 600 16th Street, Box 2280, University of California San Francisco, San Francisco, CA 94158-2517.

## Abstract

3.

The hepatic endoplasmic reticulum (ER)-anchored monotopic proteins, cytochromes P450 (P450s) are enzymes that metabolize endobiotics (physiologically active steroids and fatty acids) as well as xenobiotics including therapeutic/chemotherapeutic drugs, nutrients, carcinogens and toxins. Alterations of hepatic P450 content through synthesis, inactivation or proteolytic turnover influence their metabolic function. P450 proteolytic turnover occurs via ER-associated degradation (ERAD) involving ubiquitin (Ub)-dependent proteasomal degradation (UPD) as a major pathway. UPD critically involves P450 protein ubiquitination by E2/E3 Ub-ligase complexes. We have previously identified the ER-polytopic gp78/AMFR (autocrine motility factor receptor) as a relevant E3 in CYP3A4, CYP3A23 and CYP2E1 UPD. We now document that liver-conditional genetic ablation of gp78/AMFR in mice disrupts P450 ERAD, resulting in significant stabilization of Cyp2a5 and Cyps 2c, in addition to that of Cyps 3a and Cyp2e1. More importantly, we establish that such stabilization is of the functionally active P450 proteins, leading to corresponding significant enhancement of their drug metabolizing capacities. Our findings with clinically relevant therapeutic drugs (nicotine, coumarin, chlorzoxazone, and acetaminophen) and the prodrug (tamoxifen) as P450 substrates, reveal that P450 ERAD disruption could influence therapeutic drug response and/or toxicity, warranting serious consideration as a potential source of clinically significant drug-drug interactions (DDIs). Because gp78/AMFR is not only an E3 Ub-ligase, but also a cell-surface prometastatic oncogene that is upregulated in various malignant cancers, our finding that hepatic gp78/AMFR-knockout can enhance P450-dependent bioactivation of relevant cancer chemotherapeutic prodrugs is of therapeutic relevance and noteworthy in prospective drug design and development.

**Significance Statement:** The cell surface and ER transmembrane protein gp78/AMFR, a receptor for the prometastatic autocrine motility factor (AMF), as well as an E3 ubiquitin-ligase involved in the ERAD of not only the tumor metastatic suppressor KAI1, but also of hepatic cytochromes P450, is upregulated in various human cancers, enhancing their invasiveness, metastatic potential and poor prognosis. Liver specific gp78/AMFR genetic ablation results in functional protein stabilization of several hepatic P450s and consequently enhanced drug and prodrug metabolism, a feature that could be therapeutically exploited in the bioactivation of chemotherapeutic prodrugs, through design and development of novel short-term gp78/AMFR chemical inhibitors.

## Introduction

Luminal and membrane-bound endoplasmic reticulum (ER)-proteins incur proteolytic turnover via ubiquitin (Ub)-dependent proteasomal degradation (UPD) in a physiological process termed “ER-associated degradation (ERAD)” (Olzmann et al., 2013; Christianson and Ye, 2017; Preston and Brodsky, 2017). This vital process is critical for protein quality control required to mitigate the unfolded protein response (UPR) triggered by ER-stress and/or other cellular stresses, as well as for the normal physiological ER-protein turnover (Olzmann et al., 2013; Christianson and Ye, 2017; Preston and Brodsky, 2017). Accordingly, we have found that the hepatic monotopic ER-anchored cytochromes P450 (P450s; CYPs) incur ERAD not just upon their mechanism-based inactivation that structurally damages the proteins, but also in the course of their physiological turnover (Correia et al., 1992; Wang et al., 1999; Faouzi et al., 2007). Human hepatic P450s are responsible for the metabolism and elimination of ≈ 74% of therapeutically relevant drugs and participate in many drug-drug interactions (DDIs) and drug-related toxicities (Guengerich, 2015; Correia, 2018). The ERAD of human and rat hepatic P450s, CYP3A4, CYP3A23 and CYP2E1, involves an initial Ser/Thr-phosphorylation by protein kinases A and C (Eliasson et al. 1992, 1994; Wang et al., 2001; Wang et al., 2009, 2011, 2012, 2015), their ubiquitination via Ub-activation enzyme E1 and E2/E3 Ub-conjugation complexes (Morishima et al., 2005; Pabarcus et al., 2009; Kim et al., 2010; Wang et al., 2009, 2011, 2012), their extraction from the ER by p97/Ufd1/Npl4-AAA ATPase complex and subsequent delivery to the 26S proteasome (Liao et al., 2005; Faouzi et al., 2007; Acharya et al., 2010).

*In vitro* functional reconstitution studies (Morishima et al., 2005; Pabarcus et al., 2009; Kim et al., 2010) of E1/E2/E3-mediated CYP3A4 and CYP2E1 ubiquitination have led to the identification of UbcH5a/Hsp70/CHIP and UBC7/gp78/AMFR-complexes as two relevant E2/E3 systems in CYP3A4 and CYP2E1 ubiquitination: CHIP (Carboxy-terminus of Hsc70-interacting protein), a cytoplasmic Hsc70-cochaperone, functions with its cognate UbcH5a E2 and Hsc70/Hsp70 in substrate ubiquitination (Ballinger at al., 1999; Connell et al., 2001; Murata et al., 2001; Jiang et al., 2001). gp78/AMFR (autocrine motility factor receptor) is a polytopic, transmembrane cell surface protein (Nabi et al., 1990), as well as ER-integral protein with a cytoplasmic domain containing a RING-finger and other subdomains critical to its recruitment of its cognate E2 (UBC7 or Ube2G2) and catalytic E3-ligase-mediated ubiquitination role (Fang et al., 2001; Chen et al., 2012; Joshi et al., 2017).

Our lentiviral shRNA interference (shRNAi) analyses targeted individually against CHIP and gp78 verified their roles in CYP3A and CYP2E1 ubiquitination and ERAD in cultured rat hepatocytes (Kim et al., 2010). Thus, upon CHIP-knockdown, CYP3A was stabilized largely as the parent (55 kDa) species along with a minor CYP3A fraction consisting of its high molecular mass (HMM) ubiquitinated species. Similar stabilization of CYP3A parent species was also found upon gp78-knockdown, albeit with an appreciable fraction as its ubiquitinated species. Nevertheless, upon either E3-knockdown, the functionally active CYP3A fraction, albeit proportional to the relative amount of the stabilized parent protein, was significantly greater than that in the corresponding control (non-targeting) shRNA-treated hepatocytes (Kim et al., 2010). Furthermore, *in vitro* reconstitution studies of each CYP3A4 E2/E3-ubiquitination system, individually and in combination, revealed that although each system by itself was capable of CYP3A4-ubiquitination, this was both greatly accelerated and potentiated when both systems were present simultaneously (Wang et al., 2015). Thus, the sequential introduction of each E2/E3 in the reconstituted CYP3A4 ubiquitination system revealed that UbcH5a/CHIP most likely serves as the E3, whereas UBC7/gp78 serves as the E4 involved in the elongation of CYP3A4 polyUb chains (Wang et al., 2015).

Consistent with shRNAi findings, Cyp2e1 was also stabilized in a functionally active form upon mouse CHIP-knockout (KO) (Kim et al., 2016a). This functional Cyp2e1 stabilization was associated with increased hepatic lipid peroxidation, microvesicular steatosis, and eventually upon aging, with macrovesicular steatosis and liver injury, characteristic of the nonalcoholic steatohepatitis (NASH) syndrome (Kim et al., 2016a).

To determine whether genetic ablation of gp78, the other E3 Ub-ligase identified as relevant to hepatic CYP3A4 and CYP2E1 ERAD, had similar consequences, we generated a mouse liver conditional gp78-KO (gp78^−/−^) model. This gp78^−/−^-model enabled us not only to examine the role of gp78-mediated ubiquitination in the ERAD and consequent physiologic function of Cyp3as and Cyp2e1, but also to identify additional hepatic P450s as potential gp78-ubiquitination targets and to determine their therapeutic/toxicological consequences. Our findings are described herein.

## Materials and Methods

### Generation of a liver conditional gp78-KO mouse

Cryopreserved mouse embryos carrying a conditional *amfr*^*tma1*(Komp)Wtsi^ gene-trap knockout-first allele inserted into *amfr* intron 2 with *loxP* sites flanking exon 3 of *amfr* were generated *as described* (Skarnes et al., 2011; Bradley et al., 2012), resuscitated, cultured and transplanted into oviducts of pseudopregnant females of C57BL/6J-Tyrc-Brd; C57BL/6N background, at the UC Davis Knockout Mouse Project (KOMP) Facility. Upon weaning, the heterozygous conditional gene-trap (*cgt*) genotype of the pups was confirmed by PCR analyses. Mice aged 6 weeks were shipped to UCSF LARC Transgenic Barrier Facility, where the *loxP*-flanking sequence was deleted through mating of 8-week-old heterozygous females with transgenic male mice (C57BL/6J-background) carrying Flpe (flippase) recombinase driven by the β-actin promoter (Flpe-deleter; β-actin-Flp), and of corresponding heterozygous males with female Flpe-mice. After several intercrosses, homozygous gp78-floxed (flp/flp)-mice were obtained. To generate liver-specific gp78-KOs, flp/flp-mice were crossbred with Alb-Cre/Alb-Cre (B6.Cg-Tg(Alb-cre)21Mgn/J) mice (Postic and Magnuson, 2000). The resulting flp/+, Alb-Cre/+ mice were then crossed again with flp/flp mice to generate flp/flp, Alb-Cre/+ mice as liver-specific gp78-KOs. Mice were fed a standard chow-diet and maintained under 12 h light/dark cycle. All animal experimental protocols were approved by the UCSF/Institutional Animal Care and Use Committee (IACUC).

### Primary mouse hepatocyte culture treatment with P450 inducers

Hepatocyte isolation from male gp78+/+:Alb-Cre (WT; gp78^+/+^) and gp78−/−:Alb-Cre (KO; gp78^−/−^) mice (8 wk old) was carried out by *in situ* liver perfusion with collagenase and purification by Percoll-gradient centrifugation by the UCSF Liver Center Cell Biology Core as described previously (Han et al., 2005). Fresh primary hepatocytes (3.0 × 10^6^ cells/plate) were cultured on 60 mm Permanox plates (Thermo Fisher Scientific, Waltham, MA) coated with Type I collagen, and incubated in William’s E Medium containing 2 mM L-glutamine, Penicillin-Streptomycin, insulin-transferrin-selenium, 0.1% bovine serum albumin Fraction V, and 0.1 μM dexamethasone (DEX). Cells were allowed to attach to the plates for 6 h, before Matrigel (Matrigel^®^ Matrix, Corning Inc. New York) was overlaid. After a 2-day stabilization, cultured primary hepatocytes were treated for 3 days with the following P450-inducers: β-naphthoflavone (β-NF; 25 *μ*M; Cyp1a2), phenobarbital (PB; 1 mM; Cyps 2a, 2b and 2c), DEX (10 μM; Cyps 3a), and isoniazid (INH; 1 mM, Cyp2e1).

### Determination of P450 protein content in cultured gp78-KO mouse hepatocytes

Upon inducer-treatment, cells were harvested in cell lysis buffer (Cell Signaling Technology, Inc. Danver, MA). Lysates were cleared by centrifugation (14,000xg, 10 min, 4°C), and P450 proteins in the supernatant were subjected to Western immunoblotting (**IB**) analyses. Commercial primary antibodies were used for detecting Cyp1a2 (mouse monoclonal, Cat. No. 14719, Cell Signaling Technology, Inc.), Cyp2a (rabbit polyclonal, Cat. No. ab3570, Abcam, Cambridge, UK), and Cyp2d (rabbit polyclonal, Cat. No. GTX56286, Genetex, Inc. Irvine, CA). Purified antibodies from rabbit (Cyp2b, Cyp2c, and Cyp2e1) and from goat (Cyps 3a) sera were also used as previously described (Kim et al., 2016a, b). Immunoblots were densitometrically quantified through NIH Image J software analyses.

### Relative P450 mRNA expression in cultured WT and gp78-KO mouse hepatocyte cultures

βNF- and INH-pretreated cultured WT and gp78-KO hepatocytes were used for Cyp1a2 and Cyp2e1 mRNA level determination, respectively; PB-pretreated hepatocytes were used for monitoring Cyp2a5, 2b10, 2c29, 2c39, and 2d10 mRNA levels, whereas DEX-pretreated hepatocytes were used for monitoring Cyp3a11 and 3a13 mRNA levels. On day 4, total RNA was extracted from the cultured hepatocytes with RNeasy mini-kit (Qiagen, Hilden, Germany), and cDNA was synthesized using SuperScript IV VILO Master Mix (Thermo Fisher Scientific). Hepatic P450 mRNA expression was determined by quantitative real-time polymerase chain reaction (qRT-PCR) analyses, using SYBR Green PCR Master Mix (Applied Biosystems, Foster City, CA), and monitored by Agilent Mx3005P System (Agilent Technologies, Santa Clara, CA). Relative gene expression level of each P450 was determined by normalizing its threshold cycle (Ct) to that of GAPDH Ct. Primers used are listed (Table 1).

**Table 1.**
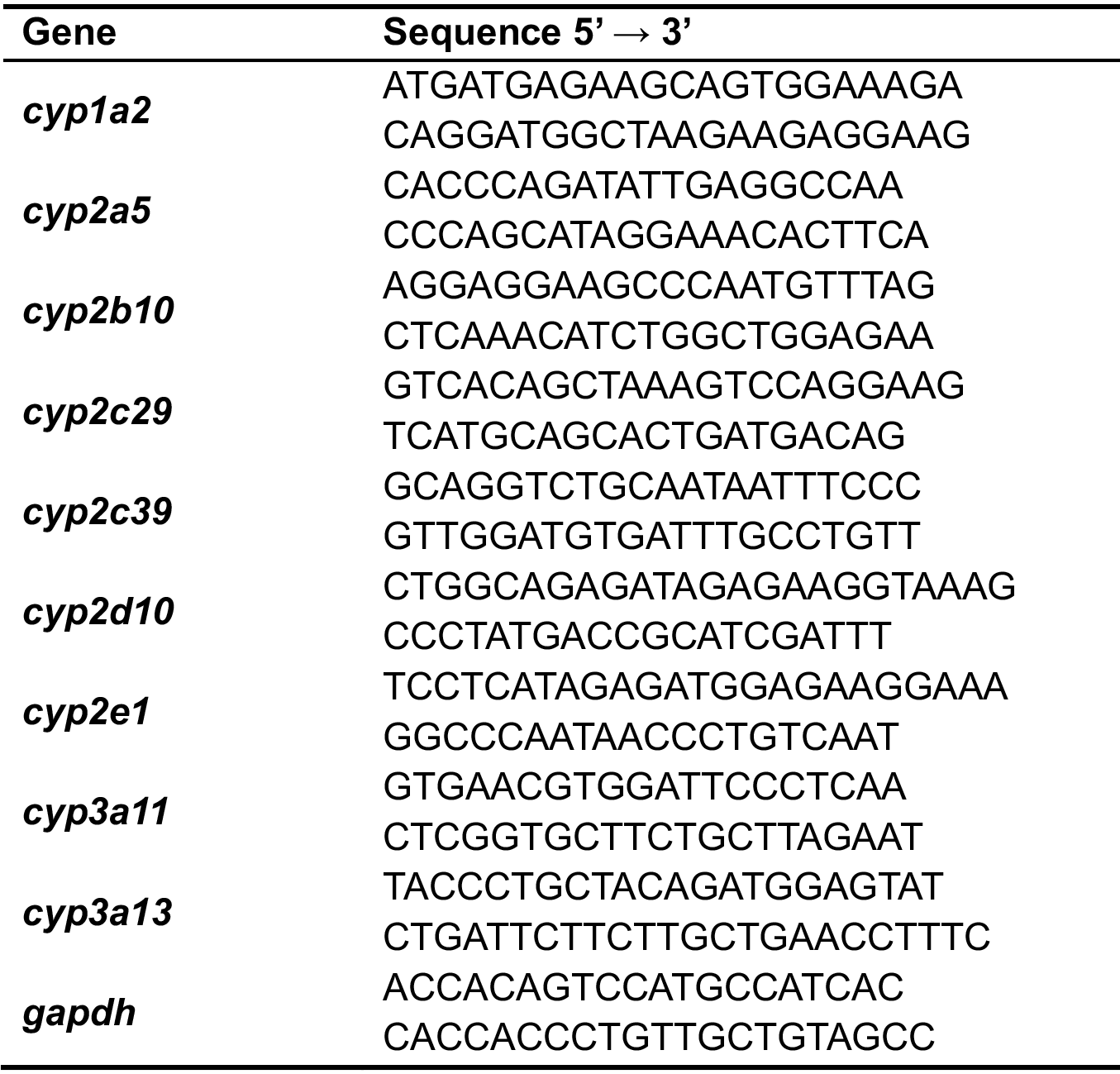
Primers used for qRT-PCR analyses of P450 mRNA expression.

### Cyp2a ubiquitination and degradation analyses

Cultured PB-pretreated hepatocytes were subjected to cycloheximide-chase analyses by treating on day 4 at 0 h with cycloheximide (CHX) 50 μg/mL with or without the proteasome inhibitor bortezomib (10 μM), or the autophagy-lysosomal degradation (ALD) inhibitors, 3-methyladenine (3-MA; 5 mM)/NH_4_Cl (50 mM) for 0, 8 and 24 h. Cells were harvested and the lysates were used for Cyp2a IB analyses.

Cyp2a5 ubiquitination analyses were conducted following P450 immunoprecipitation from the lysates (200 μg protein) of hepatocytes treated with or without bortezomib (10 μM) for 8 h. Immunoprecipitates were incubated with Cyp2a antibody at 4°C overnight. The antibody-antigen complexes were captured by incubation with Dynabeads™ protein G (Thermo Fisher Scientific) at room temperature for 2 h, and then eluted by heating at 95°C for 10 min in 2x volume of SDS-PAGE loading buffer. The ubiquitinated P450s were detected by Western IB analyses with an Ub-antibody, as described previously (Wang et al., 1999; Kim et al., 2010; Wang et al., 2015).

### Fluorescence-based screening of P450 functional markers in cultured mouse hepatocytes

P450 functional activities in cultured primary hepatocytes were determined by the method of Donato et al., (2004), as previously described (Kim et al., 2010). On day 4 after P450-inducer pretreatment, hepatocytes were incubated with P450 substrates, 3-cyano-7-ethoxycoumarin (CEC, 30 *μ*M; Cyp1a2; Cat. No. 451014. Corning Inc.), coumarin (CM, 50 *μ*M; Cyp2a5; Cat No., C4261, Sigma-Aldrich Corp., St. Louis, MO), 7-ethoxy-4-(trifluoromethyl)-coumarin (EFC, 30 *μ*M; Cyp2b10; Cat. No. T2803, Sigma-Aldrich Corp.), CEC (60 μM; Cyp2c29), dibenzylfluorescein (DBF, 10 *μ*M; Cyp2c39; Cat. No. 451740, Corning Inc.), 3-[2-(N,N-diethyl-N-methylammonium)ethyl]-7-methoxy-4-methylcoumarin (AMMC, 100 *μ*M; Cyp2d10; Cat. No. 451700, Corning Inc.), 7-methoxy-4-(trifluoromethyl)-coumarin (MFC, 10 *μ*M; Cyp2e1; Cat. No. 451740, Corning Inc.), or 7-benzyloxy-4-trifluoromethylcoumarin (BFC, 100 *μ*M; Cyp3a; Cat. No. B5057, Sigma-Aldrich Corp.) at time 0 h. The media containing CEC metabolite 3-cyano-7-hydroxycoumarin (CHC; Cat. No. 451015, Corning Inc.), CM metabolite 7-hydroxycoumarin (7-HC; Cat. No. H24003, Sigma-Aldrich Corp.), BFC, EFC and MFC metabolite 7-hydroxy-4-trifluoromethylcoumarin (HFC; Cat. No. 451731, Corning Inc.), or DBF metabolite fluorescein (Cat. No. 46955, Sigma-Aldrich Corp.), along with their glucuronide and sulfate conjugates formed during the assay were collected at defined time points. AMMC metabolite, AMHC was commercially unavailable, and thus 3-[2-(diethylamino)ethyl]-7-hydroxy-4-methylcoumarin (AHMC; Cat. No. 188611, Sigma-Aldrich Corp.) was used instead as previously described for the fluorescence quantification (Donato et al. 2004). The conjugates were hydrolyzed by incubation of the media with glucuronidase/arylsulfatase (150 Fishman units/ml and 1200 Roy units/ml, respectively) for 2 h at 37°C. The medium concentrations of CEC, 7-HC, HFC, fluorescein and AHMC were fluorometrically monitored at the wavelength [excitation (nm)/emission (nm)] of 408/455, 355/460, 410/510, 485/538, and 390/460, respectively, using the commercially purchased metabolites as authentic standards.

### Nicotine metabolism in cultured mouse hepatocytes

WT and gp78-KO hepatocytes were PB-pretreated as described above. On day 4, (−)-Nicotine (500 *μ*M) was added at time 0 h and the cells and media were collected at 2, 4, 8, and 24 h, and incubated with glucuronidase/arylsulfatase for 2 h at 37°C. The reaction was stopped by the addition of ice-cold 30% perchloric acid, and the precipitated proteins were sedimented by centrifugation (15,000xg, 10 min, 4°C). Nicotine metabolites (cotinine and *trans*-3’-hydroxycotinine, nornicotine, norcotinine, nicotine-N-oxide and cotinine-N-oxide) were extracted from the supernatant and quantified by LC-MS/MS analyses as described for cotinine and trans-3’-hydroxycotinine (Jacob et al., 2011), with minor modifications for the additional analytes (Benowitz et al., 2010, Wang et al., 2018a). The metabolites were found largely in the media, with relatively much smaller levels detected in the cell lysates.

### Tamoxifen metabolism in cultured mouse hepatocytes

Cyp3a was induced by DEX-pretreatment of primary hepatocytes as described above. On day 4, cultured hepatocytes were treated with tamoxifen (40 *μ*M) for 2, 4 and 8 h. The concentrations of tamoxifen and its metabolites, 4-hydroxytamoxifen, N-desmethyltamoxifen, and endoxifen in the media were determined using a HPLC-MS (mass spectrometric) assay (Dahmane et al., 2010; Binkhorst et al., 2011). Toremifen was employed as an internal standard, and spiked into the media at 1 *μ*M. The medium was incubated with glucuronidase/arylsulfatase for 2 h at 37°C and the reactions terminated by the addition of 2X volume of ice-cold acetonitrile containing 0.1% formic acid. Precipitated proteins were removed by sedimentation at 10,000xg for 10 min at 4°C, and an aliquot (300 *μ*L) of the supernatant was mixed with 50 mM glycine buffer (pH 11.5; 700 *μ*L). An aliquot (50 *μ*L) of this mixture was injected into the HPLC system (Waters Separation Module 2695; Waters Corporation, Milford, MA), equipped with a C18 reversed phase column (100 × 2.1 mm, 3 *μ*m, Hypersil Gold aQ; Thermo Fisher Scientific), maintained at a temperature of 30°C. An aqueous solution containing 0.1% formic acid was used as the mobile phase A, and acetonitrile containing 0.1% formic acid was used as the mobile phase B. Mobile phases A (30%) and B (70%) were applied isocratically at a flow rate of 0.2 mL/min. The peaks were detected at 254 nm by Waters dual absorbance detector 2487 (Waters Corporation). The retention times of endoxifen, 4-hydroxytamoxifen, toremifen, N-desmethyltamoxifen, and tamoxifen were 9.26, 11.16, 22.17, 23.77, and 26.92 min, respectively. The HPLC system was connected to a Waters Micromass ZQ single quadrupole mass analyzer (Waters Micromass Corporation, Manchester, U.K.), equipped with an electrospray ionization probe operated in the positive ionization mode. The ion-spray and cone voltage were 3 kV and 45 V, respectively. The protonated molecular ion [M+H]^+^ masses of N-desmethyltamoxifen, tamoxifen, endoxifen, 4-hydroxytamoxifen, and toremifen were found to be 358.2, 372.2, 374.2, 388.2, and 406.2 amu, respectively.

### Chlorzoxazone metabolism in cultured mouse hepatocytes

Cultured primary WT and KO mouse hepatocytes were INH-pretreated to induce Cyp2e1, as described above. On day 4, at time 0 h, chlorzoxazone (400 *μ*M) was added to the hepatocytes for 3 h. The 6-hydroxychlorzoxazone (6-OH CZX) formed by Cyp2e1 was monitored by HPLC as described (Girre et al., 1994) with a slight modification. The collected medium was first incubated with glucuronidase/arylsulfatase for 2 h at 37°C, and the reaction stopped with 40% perchloric acid. After sedimentation of the precipitated protein, the supernatant was mixed with 1.5X volume of ethyl acetate by vigorous vortex mixing for 20 min. The upper organic phase was collected and evaporated at 40°C. The resulting residue was reconstituted with 100 *μ*L of mobile phase, acetonitrile and 0.5% acetic acid (30:70, v:v, pH 3), and a 50 *μ*L-aliquot was injected into a Varian Prostar HPLC (Varian Inc., Palo Alto, CA) system, equipped with a Grace Prosphere 100 C18, 5 *μ*m, 4.6 × 150 mm column (Thermo Fisher Scientific) for metabolite separation at a flow rate of 1.1 mL/min. The peak of 6-HO-CZX (retention time of 7.6 min) was detected at 287 nm with a Varian Prostar 335 UV detector (Varian Inc.).

### Acetaminophen (APAP)-induced hepatotoxicity

Cultured mouse hepatocytes pretreated with INH (1 mM) for 3 d were used for the APAP toxicity assay. On day 4, hepatocytes were incubated with APAP (1 or 5 mM) and the medium was collected at 2, 4, 8, and 24 h. The medium alanine aminotransferase (ALT) activity was quantified using the ALT assay kit (MAK052, Sigma-Aldrich Corp.).

## Results

### Verification of the liver conditional gp78-KO mouse genotype

The cgt-targeting strategy employed is shown (*Fig. 1A*). This strategy is different from that reported previously (Liu et al., 2012), in that the cgt-cassette employed and the targeted exons (3 vs 5-8) were different, and it also employed resuscitated cryopreserved embryos rather than genetically engineered embryonic stem cells as the starting material. Verification of the genomic DNA by tail-clip genotyping via PCR analyses of wild-type (WT; gp78^+/+^) flp/flp-mice, cgt/+, gp78^+/−^ and gp78^−/−^-mice, respectively, is shown (*Fig. 1B, C*). Parallel IB analyses of liver and brain tissue lysates from WT and gp78^−/−^-mice provided unequivocal evidence of liver conditional gp78-KO (*Fig. 1D*).

**Fig. 1.**
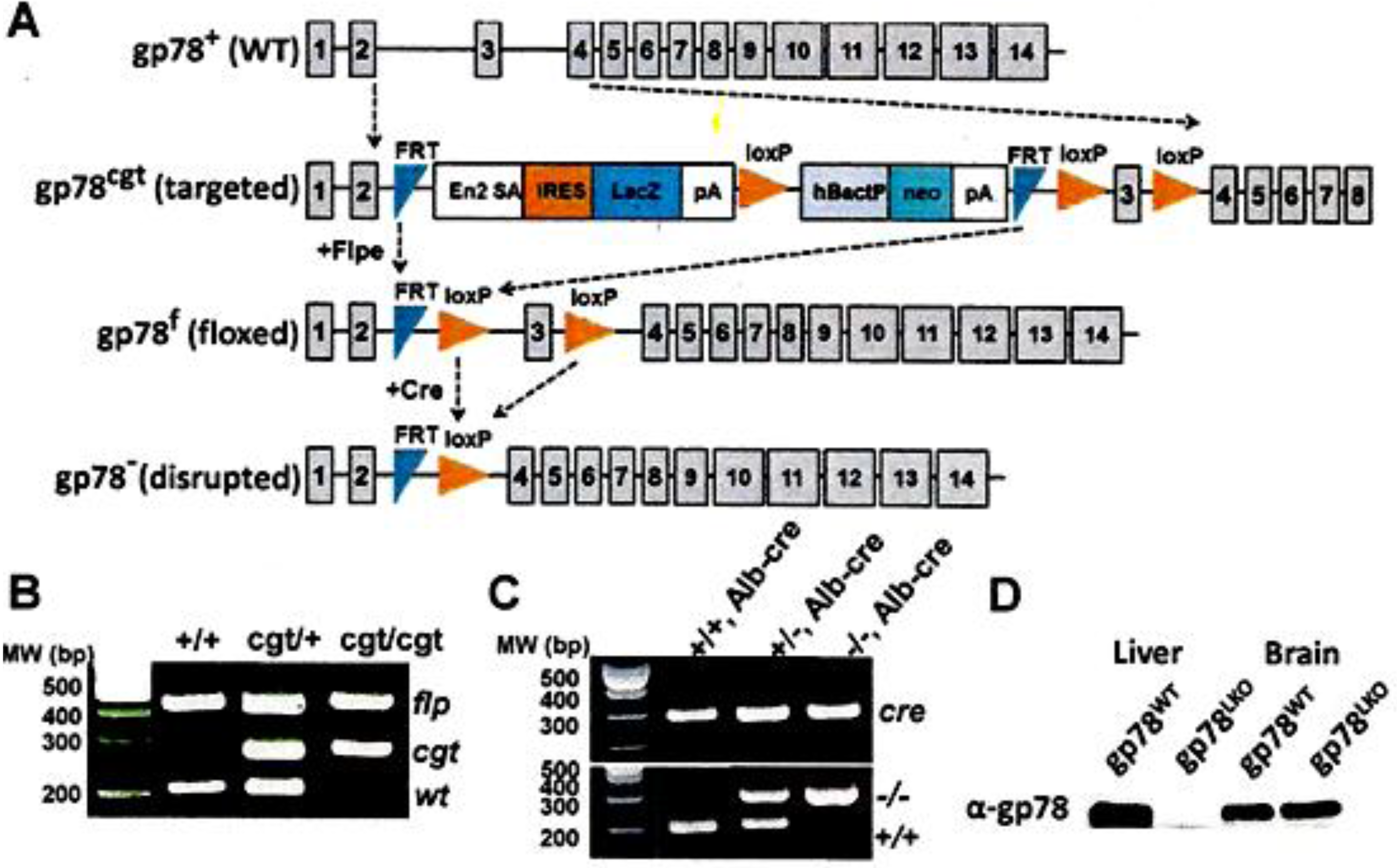
Strategy for generating the liver conditional gp78/AMFR-KO mouse model and corresponding validation: A. Conditional gene targeting (cgt) strategy to develop gp78-floxed mice and subsequently, liver conditional gp78-KO mice. B, C. PCR genotype analyses of genomic DNA isolated via tail-clipping of mice at various stages of the strategy. D. IB analyses of liver and brain lysates from WT and KO-mice with a rabbit anti-mouse gp78-antibody.

### Identification of hepatic P450s as gp78-targets

Consistent with our gp78-shRNAi findings in cultured rat hepatocytes, **IB** analyses of lysates from cultured mouse hepatocytes verified that parent Cyps 3a and 2e1 were stabilized upon gp78-KO (gp78^−/−^) relative to those in the WT (Fig. 2). Corresponding diagnostic functional probes revealed that this stabilization was largely of the functionally active forms (Fig. 3). Similar conclusions were also drawn for Cyps 2c (Figs. 2 & 3). The Cyp1a2 increase in gp78^−/−^-hepatocytes relative to the WT-hepatocytes on the other hand, may be partially due to transcriptional induction (Fig 4). By contrast, the relative hepatic content and corresponding functional activity of Cyp2d10 were unaffected, whereas those of Cyp2b10 were even decreased upon gp78-KO (Figs. 2 & 3). Surprisingly, a marked stabilization of Cyp2a5 was observed in gp78^−/−^- over gp78^+/+^-hepatocytes (Fig. 2), with a correspondingly marked relative increase of coumarin-7-hydroxylase in gp78^−/−^-over gp78^+/+^-hepatocytes (Fig. 3). Thus, in addition to Cyps 3a and Cyp2e1, these studies identified Cyp2a5 and Cyps 2c as gp78-targets, with their diagnostic functional probes attesting to their corresponding statistically significant functional increases.

**Fig. 2.**
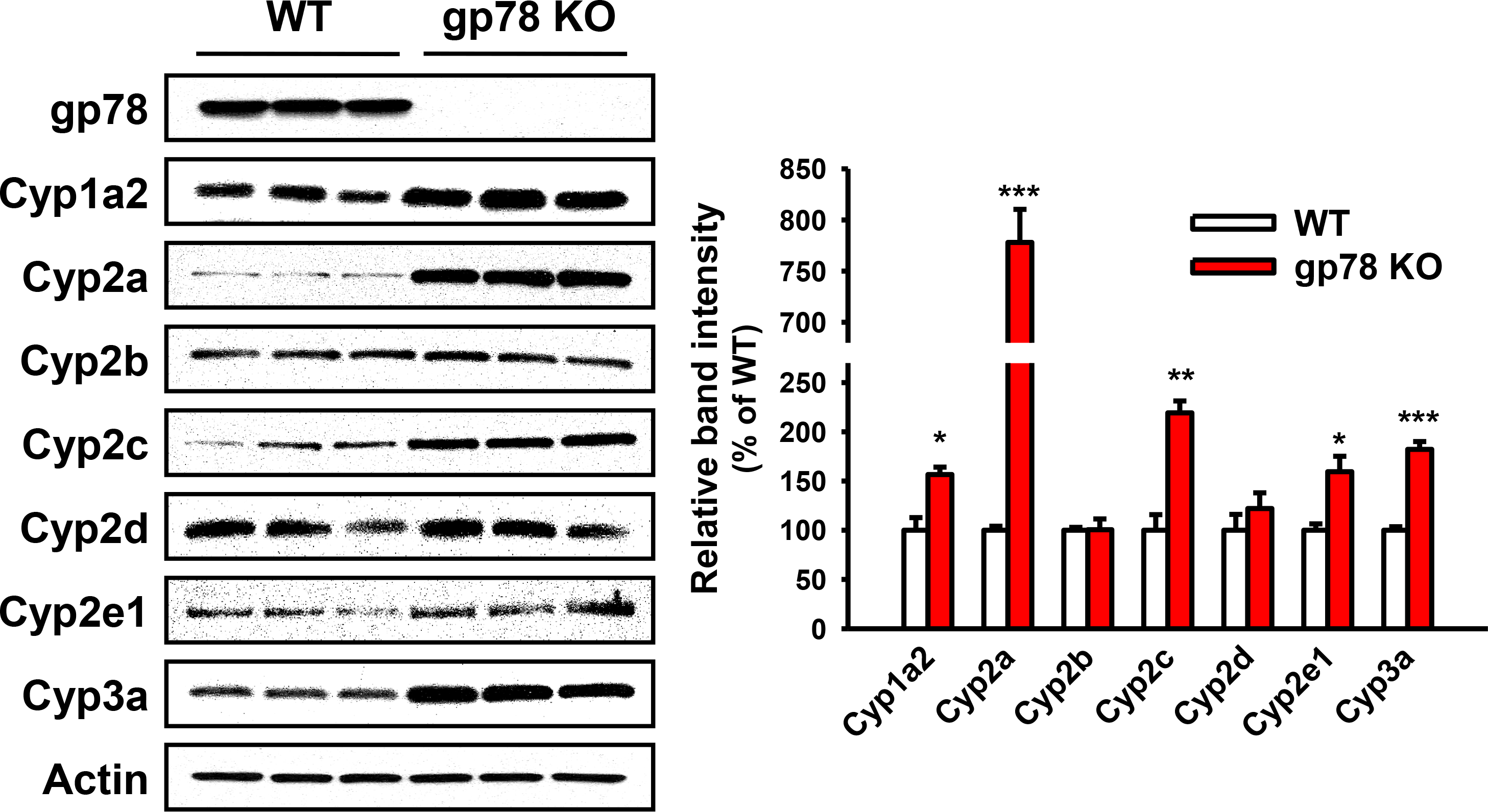
IB analyses of hepatic P450s from WT and gp78-KO mice: A. IB analyses of a representative mouse sample. B. Densitometric quantification of the ≈ 50-55 kDa P450 band from hepatocytes isolated from 3 individual WT and gp78-KO mice, cultured and pretreated with P450-inducers. See Materials & Methods for details. Values (Mean ± SD) of 3 individual experiments. Statistical significance determined by Student’s t-test, *,**,\*\*\**p* < 0.05, 0.01, and 0.001 vs. WT, respectively.

**Fig. 3.**
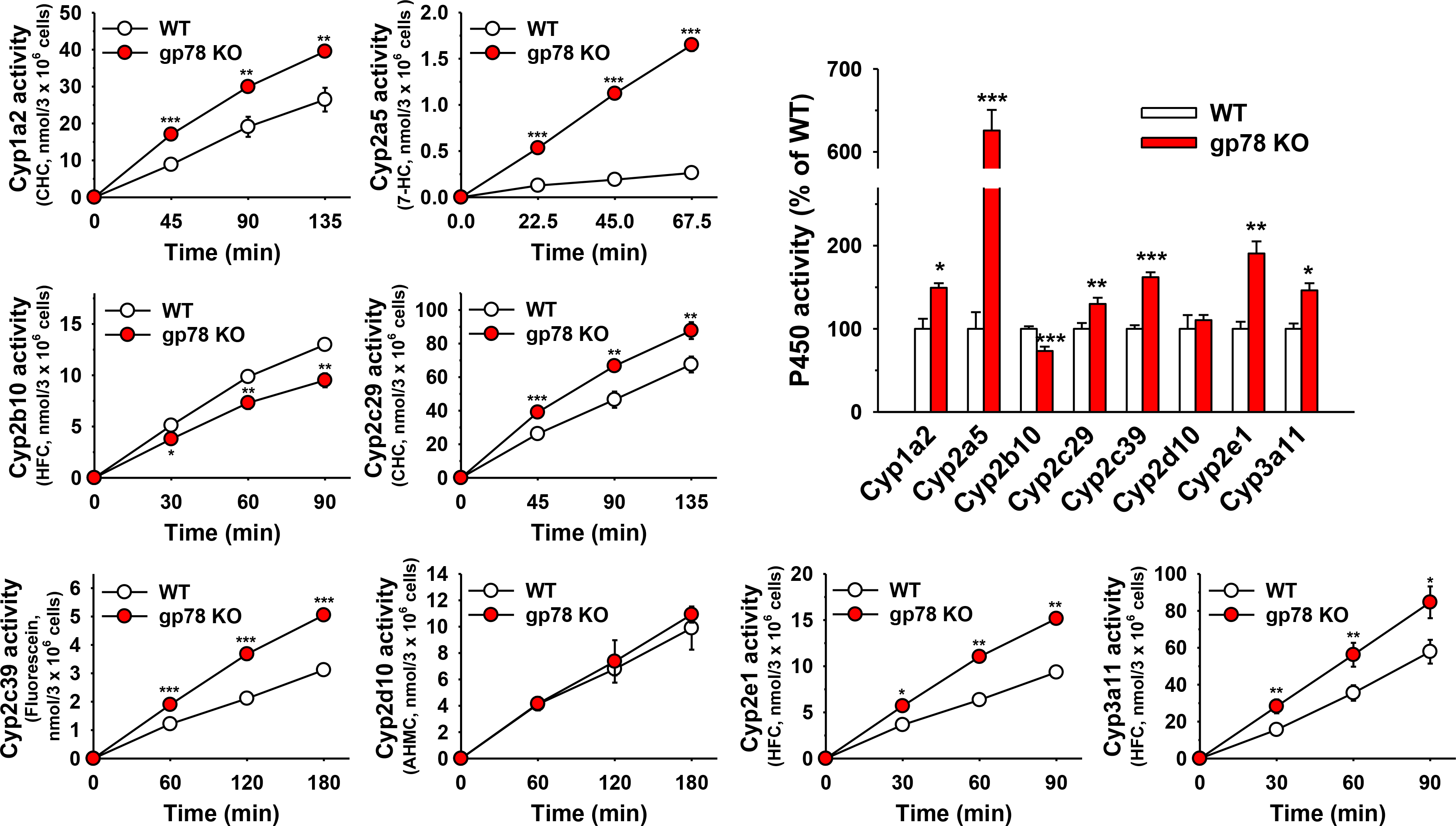
P450 functional screening using diagnostic fluorescent probes: Hepatocytes from 3 individual WT and gp78-KO mice were cultured, pretreated with P450-inducers, and then incubated with individual diagnostic functional probes at time 0 h. See Materials & Methods for details. Values (Mean ± SD) of 3 individual experiments. Statistical significance determined by Student’s t-test, *,**,\*\*\**p*< 0.05, 0.01, and 0.001 vs. WT, respectively.

**Fig. 4.**
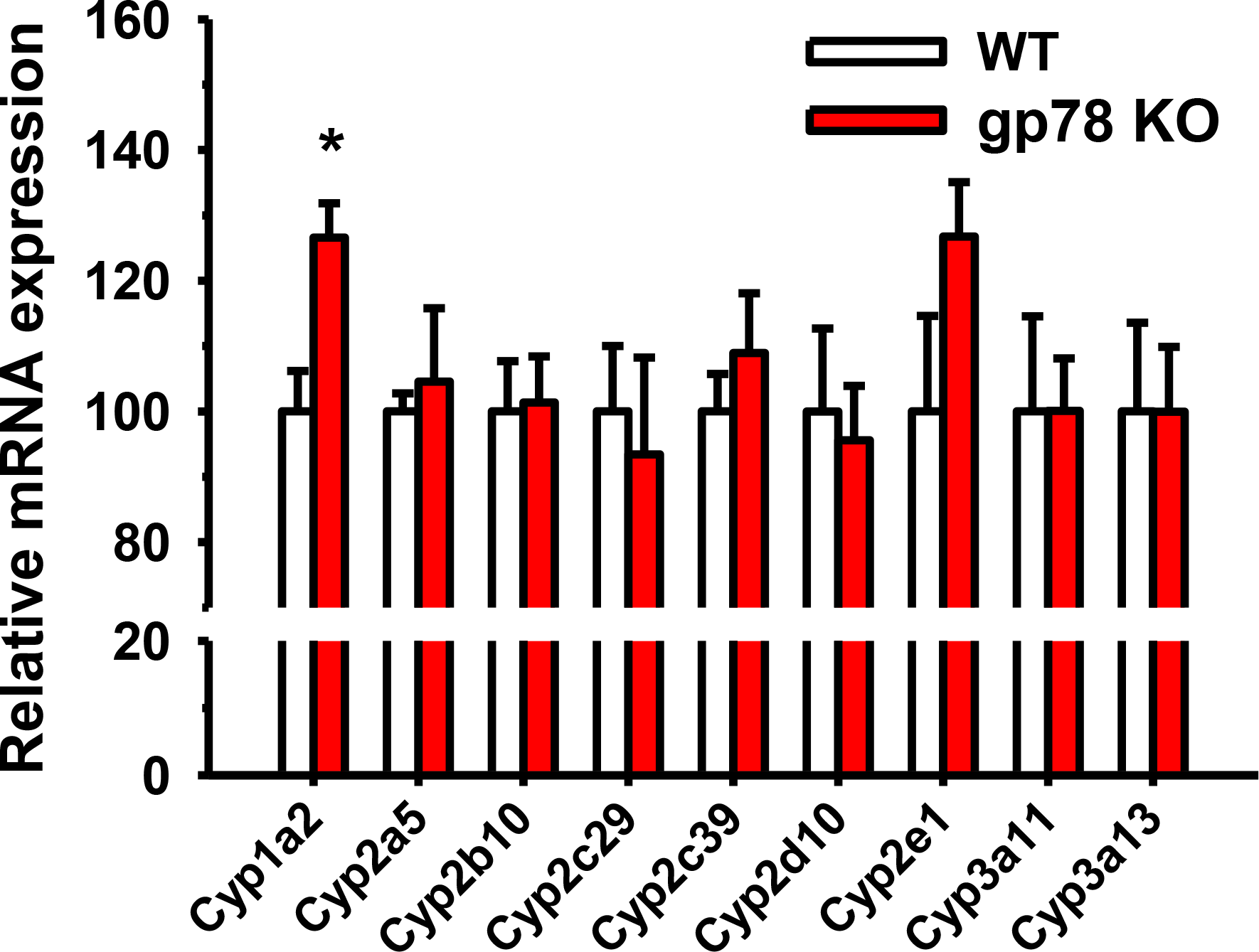
Relative P450 mRNA expression in cultured WT and gp78-KO mouse hepatocytes: Total RNA was extracted on day 4 from cultured hepatocytes, untreated or pretreated with P450 inducers for 3 days. Hepatic P450 mRNA expression was determined by qRT-PCR analyses as detailed in Materials & Methods. Values (Mean ± SD) are derived from 3 individual cultures pretreated with P450 inducers as detailed. Statistical significance determined by Student’s t-test, \**p*< 0.05 vs. WT.

### Functional P450-enhancement is due to corresponding P450 protein stabilization upon gp78-KO

We have previously reported that the CYP3A stabilization upon lentiviral shRNAi-elicited gp78-knockdown in cultured rat hepatocytes is due to its protein stabilization rather than increased CYP3A mRNA expression (Kim et al., 2010). Relative hepatic P450 mRNA expression analyses by qRT-PCR analyses of total RNA isolated from gp78^−/−^- and gp78^+/+^-hepatocytes after each P450 inducer-pretreatment revealed that the observed relative P450 content and functional increases (except in the case of Cyp1a2), were not due to their transcriptional mRNA induction (Fig. 4).

Cyp2a5-immunoprecipitation from PB-pretreated gp78^−/−^- and gp78^+/+^-hepatocytes and protein ubiquitination analyses, directly verified that Cyp2a5 was primarily a gp78-target, as its ubiquitinated level was markedly reduced upon gp78-KO, and bortezomib-mediated inhibition of its proteasomal degradation failed to enhance it (Fig. 5A). Furthermore, cycloheximide-chase analyses of gp78^−/−^- and gp78^+/+^-hepatocytes coupled with concurrent treatment with either the proteasome inhibitor bortezomib, or the ALD inhibitors (3-MA/NH_4_Cl) revealed that gp78-mediated ubiquitination targeted Cyp2a5 to both UPD and ALD (Fig. 5B).

**Fig. 5.**
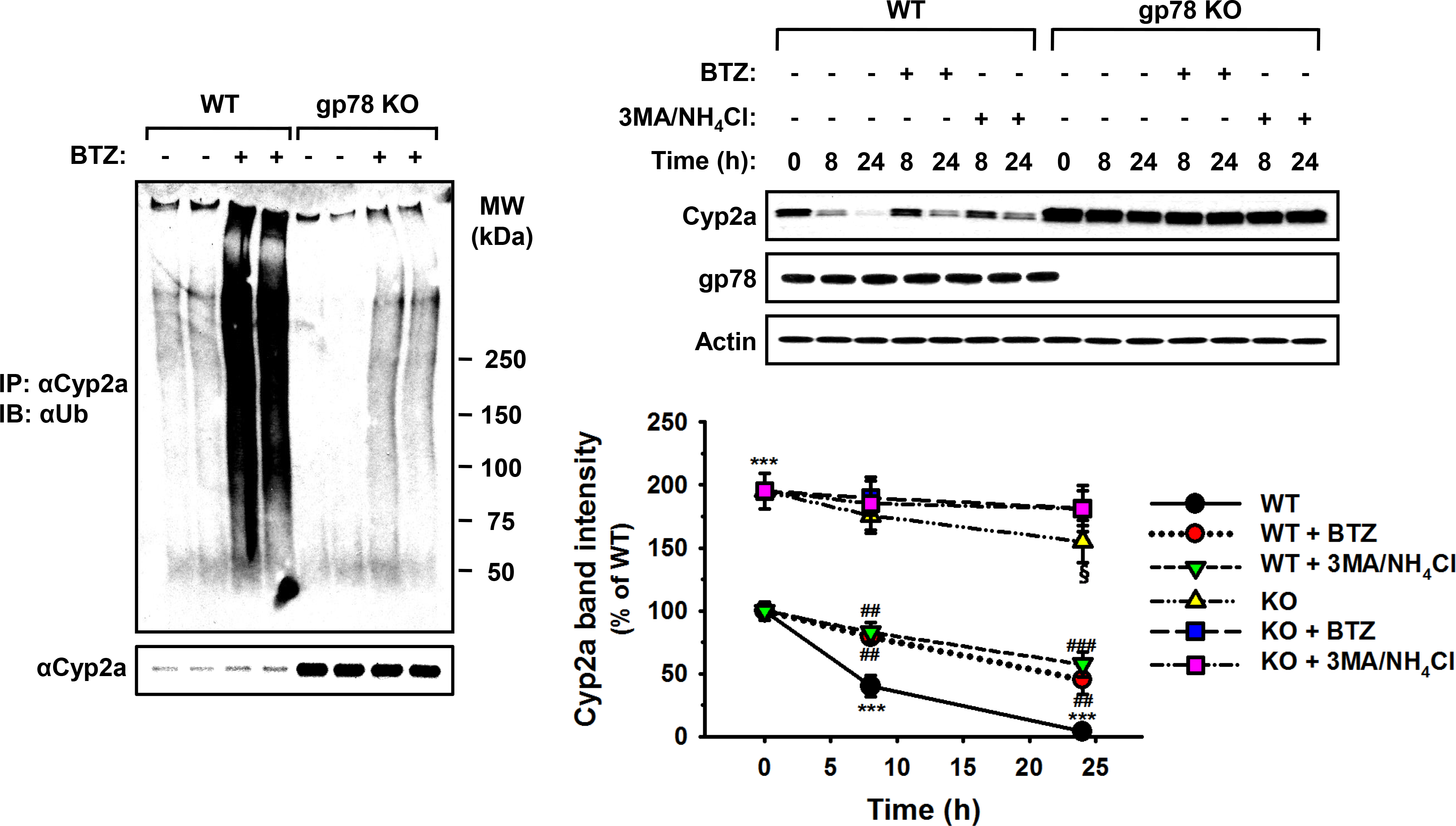
Hepatic Cyp2a5 UPD and ALD: A major role of gp78 in Cyp2a5 ubiquitination. A. Cycloheximide (CHX)-chase analyses of 3 individual, PB-pretreated WT and gp78-KO mouse hepatocyte cultures treated on day 4 with or without the proteasomal inhibitor bortezomib (BTZ) or the ALD-inhibitors 3-MA/NH_4_Cl for 0, 8 and 24 h before cell harvesting. Cell lysates were immunoblotted for Cyp2a or gp78 with actin as a loading control. B. Densitometric quantification (Mean ± SD) of 3 individual experiments. Statistical significance determined by Student’s t-test, *** *p*< 0.001 vs WT at 0 h; ^##, ###^ *p* < 0.01, 0.001 vs WT+ CHX; ^@^*p*< 0.05 vs KO at 0 h.

### Pharmacological relevance of hepatic P450 stabilization upon gp78-KO

The above functional analyses with the exception of coumarin, employed readily assayable fluorescent diagnostic probes rather than therapeutic and/or pharmacologically active drugs as substrates. To determine the potential therapeutic relevance of CYP2e1-stabilization upon gp78-KO, we examined the Cyp2e1-dependent 6-hydroxylation of chlorzoxazone, a clinically prescribed skeletal muscle relaxant and an established Cyp2e1 functional probe, in gp78^−/−^- and gp78^+/+^-hepatocytes (Fig. 6). A statistically significant relative increase was detected in 6-OH-chlorzoxazone formation in gp78^−/−^- over gp78^+/+^-hepatocytes (Fig. 6), comparable to that observed with its other diagnostic Cyp2e1 probe (MFC) (Fig. 3), thus underscoring the potential therapeutic significance of these findings.

Cyp2e1 largely, and Cyps 3a to a lesser extent, are also involved in the metabolism of the clinically prescribed non-steroid analgesic acetaminophen (APAP) to its reactive nucleophilic species N-acetylparaquinoneimine (NAPQI) (Lee et al., 1996; Wolf et al., 2007). This pathway becomes increasingly hepatotoxigenic when liver concentrations of ingested APAP exceed the detoxification capability of its hepatic glucuronidation and sulfation pathways (Gillette et al., 1981; Nelson, 1990). More of the parent drug is then shunted into the reactive NAPQI-pathway with consequent liver damage that can be clinically fatal (Nelson, 1990). To determine the potential hepatotoxicological relevance of Cyp2e1 functional stabilization upon gp78-KO, we examined INH-pretreated gp78^−/−^- and gp78^+/+^-hepatocytes treated with APAP at two different concentrations: 5 mM, the concentration routinely employed to trigger APAP-induced liver damage (Burcham and Harman, 1991; Bajt et al., 2004), and 1 mM, one-fifth the toxic concentration, over 24 h of culture. Although as expected, the 5 mM concentration led to significantly increased injury to gp78^+/+^-hepatocytes, as monitored by the extracellular spillage into the cell medium of the liver cytoplasm-specific ALT enzyme, this was significantly greater in gp78^−/−^-hepatocytes (Fig. 7). At the 1 mM-concentration however, while APAP produced a mild extracellular ALT-rise in gp78^+/+^-hepatocytes, this was considerably accelerated and pronouncedly greater in gp78^−/−^-hepatocytes (Fig. 7). These findings thus attest to the relevance of P450 stabilization upon gp78-KO to drugs whose hepatotoxigenic potential largely depends on their Cyp2e1-mediated bioactivation.

**Fig. 6.**
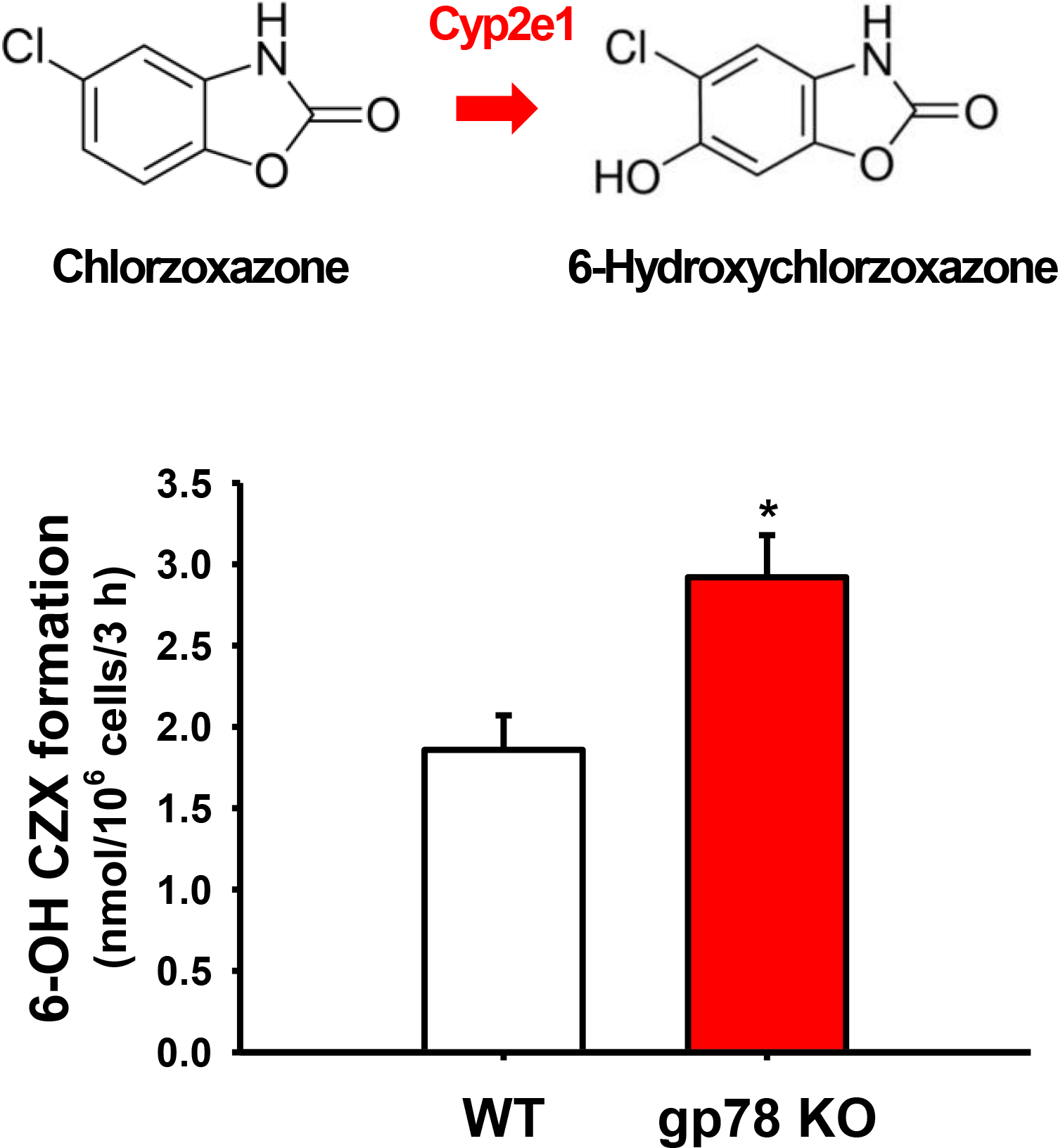
Relative chlorzoxazone metabolism in cultured WT and gp78-KO mouse hepatocytes: INH-pretreated hepatocytes from 3 individual WT and gp78-KO mice were cultured, and on day 4 incubated with chlorzoxazone for 3 h and 6-hydroxychlorzoxazone assayed as detailed in Materials & Methods. Values (Mean ± SD) of 3 individual experiments. Statistical significance determined by Student’s t-test, \**p*< 0.05 vs. WT.

**Fig. 7.**
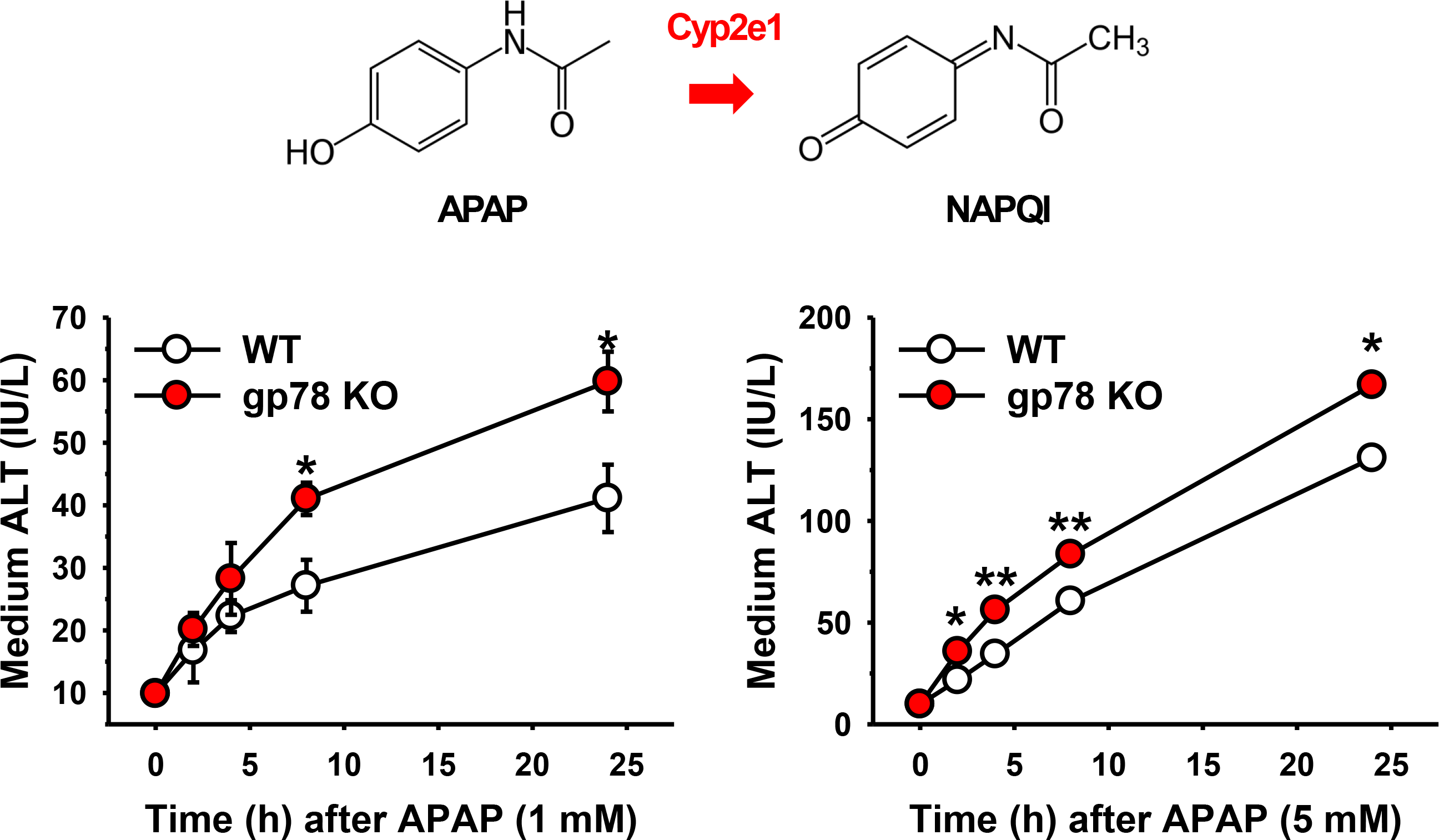
Relative APAP-elicited liver injury in cultured WT and gp78-KO mouse hepatocytes: Three individual WT and gp78-KO mouse hepatocyte cultures were INH-pretreated, and on day 4 incubated with APAP (1 or 5 mM) for 0-24 h. Aliquots of media were taken at the time points indicated and assayed for ALT activity as detailed in Materials & Methods. Values (Mean ± SD) of 3 individual cultures. Statistical significance determined by Student’s t-test, \**p*< 0.05 vs. WT.

Furthermore, to determine the pharmacological relevance of the relatively marked functional Cyp2a5-stabilization, we also determined its effects on the metabolism of its prototype drug substrate nicotine, a pharmacologically active drug (Fig. 8). As expected, nicotine was metabolized to cotinine (via the nicotine-iminium ion and subsequent oxidation by aldehyde oxidase), nornicotine, norcotinine, *trans*-3’-hydroxycotinine, nicotine-N-oxide and cotinine N-oxide (Cashman et al., 1992; Nakajima et al., 1996; Hukkanen et al., 2005; Siu et al., 2006; Zhou et al., 2010). While the nicotine N-oxide is reportedly generated by flavin-monooxygenase (FMO3; Cashman et al., 1992), the other 4 metabolites are generated by various P450s, of which Cyp2a5 is the major catalyst (Nakajima et al., 1996; Siu et al., 2006; Zhou et al., 2010). On the other hand, *trans*-3’-hydroxycotinine is selectively generated by CYP2A6, a Cyp2a5 ortholog (Nakajima et al., 1996; Siu et al., 2006; Zhou et al., 2010). Our results revealed that indeed, from 2 h onwards, consistent with the marked increase in Cyp2a5, there is a statistically significant, accelerated increase in cotinine formation, with a 2-fold increase in *trans*-3’-hydroxycotinine formation detected at 24 h in gp78^−/−^- over gp78^+/+^-hepatocytes (Fig. 8). Thus, not quite as high as the marked functional fold-increases observed with the diagnostic Cyp2a functional marker (coumarin) (Fig. 3), but significant nonetheless. One issue with monitoring secondary drug metabolism in isolated cultured hepatocytes in general, is that the primary metabolite (cotinine) spills out into the cell medium almost as quickly as it is generated, thus not allowing sufficient time for its subsequent Cyp2a5-selective *trans*-3’-hydroxycotinine conversion. Thus, only when sufficient cotinine accumulates intracellularly with time (24 h), is the *trans*-3’-hydroxycotinine conversion detectable^1^ (Fig. 8). The other is that nicotine itself may incur enhanced substrate competition by other P450s that not only are known to metabolize it (Zhou et al., 2010), but are also increased upon gp78-KO, which is not the case with the relatively more selective Cyp2a5 diagnostic probe. Nevertheless, these findings indicate that gp78-KO can significantly increase the metabolism of pharmacologically active drugs by enhancing hepatic Cyp2a5 levels through protein stabilization.

**Fig. 8.**
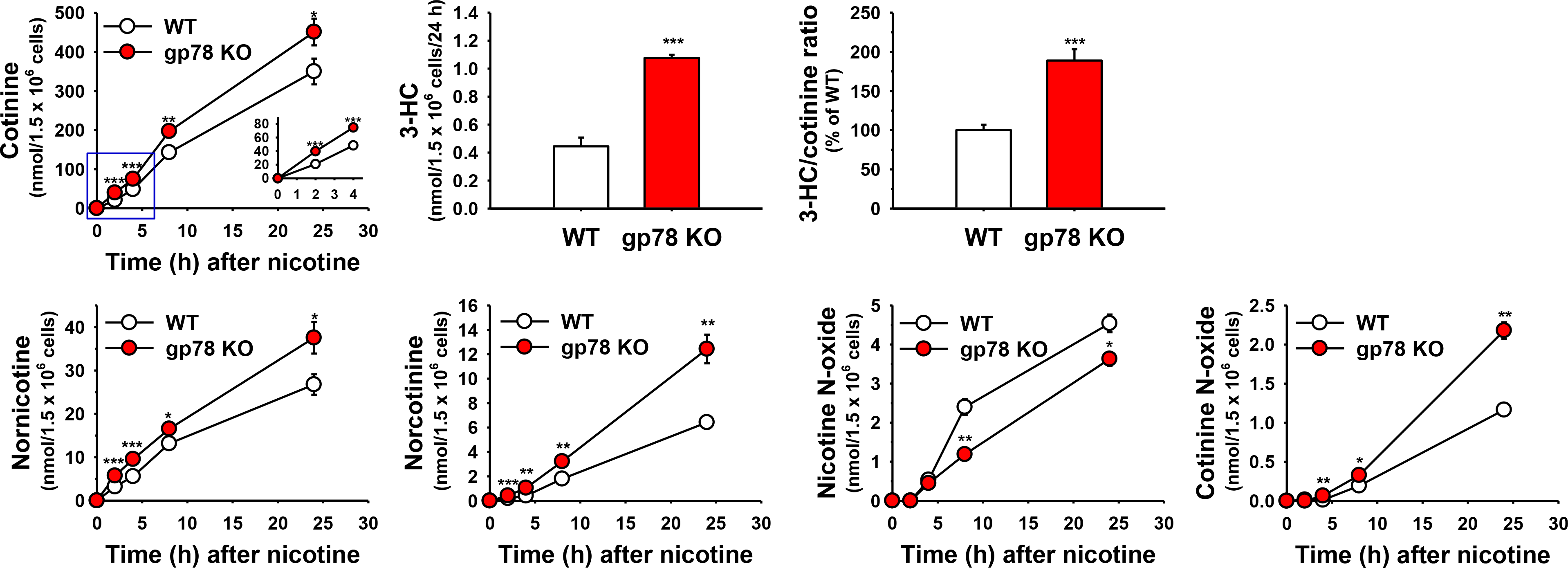
Relative nicotine metabolism in cultured WT and gp78-KO mouse hepatocytes: PB-pretreated hepatocytes from WT and gp78-KO mice were cultured, and on day 4 incubated with nicotine for 0-24 h. Aliquots of media were taken at the time points indicated and assayed as detailed in Materials & Methods. Values (Mean ± SD) of 3 individual cultures. Statistical significance determined by Student’s t-test, \**p*< 0.05 vs. WT.

Pharmacologically relevant drugs often involve multiple hepatic P450s, each catalyzing a different step of their biotransformation pathway. Such biotransformation is relevant not only to their elimination, DDIs and toxicity, but also often to their pharmacological response through prodrug metabolic bioactivation. We therefore sought to determine the relevance of gp78-KO to the bioactivation of the estrogen receptor-positive breast cancer therapeutic prodrug, tamoxifen. Tamoxifen bioactivation involves two concurrent alternate sequential metabolic routes (Fig. 9; Jacolot et al., 1991; Desta et al., 2004; Goetz et al., 2007): It can be first converted to 4-hydroxytamoxifen by Cyp2d, which is then N-demethylated by Cyps 3a to generate endoxifen. Alternatively, it could be first N-demethylated by Cyps 3a to N-desmethyltamoxifen, which is then 4-hydroxylated by Cyp2d to generate endoxifen (Jacolot et al., 1991; Desta et al., 2004). Both routes thus generate endoxifen, the ultimately desired bioactive chemotherapeutic entity. Our findings reveal that upon gp78-KO, although the Cyp2d-dependent 4-hydroxylation step remains unchanged, consistent with unaltered hepatic Cyp2d levels, the bioactivation of tamoxifen to endotoxifen is nevertheless significantly increased consistent with functional Cyp3a stabilization, and would be therapeutically relevant (Fig. 9).

**Fig. 9.**
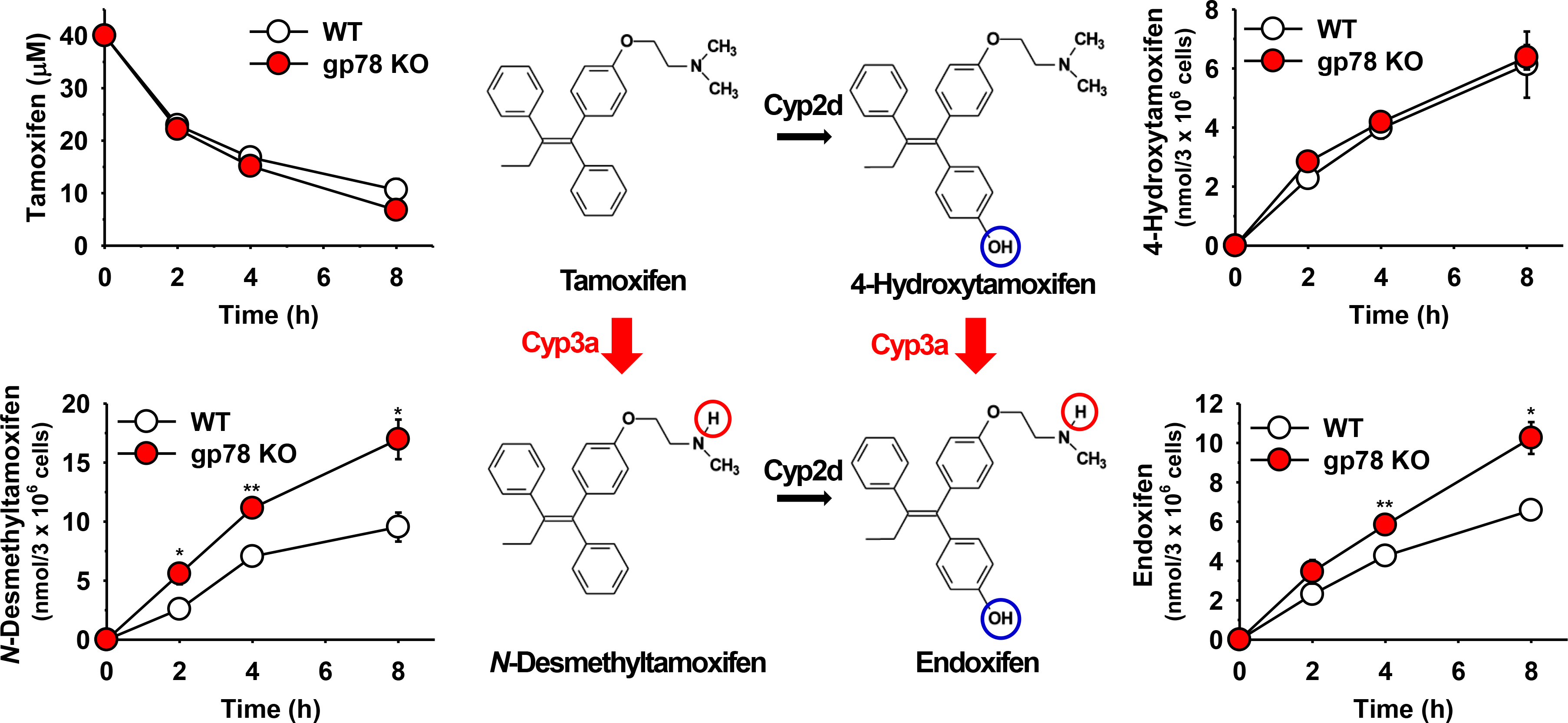
Relative tamoxifen metabolism in cultured WT and gp78-KO mouse hepatocytes: Three individual WT and gp78-KO mouse hepatocyte cultures were DEX-pretreated and on day 4 incubated with tamoxifen (40 μM) for 0-8 h. Aliquots of cell media were taken at the time points indicated and assayed as detailed in Materials & Methods. Values (Mean ± SD) of 3 individual experiments. Statistical significance determined by Student’s t-test, \**p*< 0.05 or ** *p*< 0.01 vs. WT.

## Discussion

We document herein for the first time that hepatic Cyp2a5 and Cyps 2c, in addition to previously established Cyps 3a and Cyp2e1, are stabilized upon liver conditional gp78/AMFR-KO. Furthermore, we provide concrete novel evidence that Cyp2a5 not only is primarily an ubiquitination target of gp78 E3-ligase, but also despite the preferential gp78 mode of K48-linked rather than K63-linked ubiquitination (Li et al., 2007), Cyp2a5 is targeted to both UPD and ALD. Based on our findings of their human CYP2C9 ortholog, Cyps 2c are quite likely also gp78-ubiquitination substrates (Kim et al. 2016b). Thus, the observed hepatic P450 stabilization upon disruption of their UPD through gp78-KO provides additional clinically relevant examples of eukaryotic protein induction via protein stabilization (Schimke and Doyle, 1970).

The mouse hepatic P450s stabilized upon gp78-KO, have corresponding human orthologs that are involved not just in the metabolism and elimination of many clinically prescribed drugs, dietary agents, and environmental xenobiotics, but also in their biotransformation either to more pharmacologically active or even more toxic/carcinogenic/mutagenic species (Guengerich, 2015; Correia, 2018). Employing pharmacological agents with therapeutic and/or hepatotoxigenic potential, we document herein that such hepatic P450 functional stabilization is likely of clinical relevance. Accordingly, most therapeutic agents, identified as specific substrates of a given hepatic P450 that is functionally stabilized upon gp78-KO, would experience enhanced clearance and accelerated termination of their pharmacological action. Their therapeutic responses would be consequently curtailed much akin to the impaired drug responses observed upon co-ingestion of therapeutic or dietary P450 inducers. For drugs such as nicotine metabolized predominantly by CYP2A6, its functional stabilization while accelerating nicotine clearance, may also lead to enhanced nicotine-craving in chronic cigarette-smokers, and thus to an altered behavioral response. Notably, given the major CYP2A6-role in the bioactivation of tobacco nitrosamine carcinogens, including the tobacco-specific lung carcinogen 4-(methylnitrosamino)-1-(3-pyridyl)-1-butanone (NNK) and other toxic nitrosamines (Zhou et al., 2012; Zhu et al., 2013; Yuan et al., 2017), such carcinogenecity would also be enhanced.

Importantly, because many of these P450s are also involved in the metabolism of drugs (i.e. cancer chemotherapeutic agents) with narrow therapeutic indices, even minor oscillations in hepatic P450 content may translate into therapeutically undesirable losses of their pharmacological efficacy on the one hand, and toxicity on the other, as underscored previously (Peer et al, 2011), in support of the clinical relevance of our CHIP- and gp78-shRNAi findings (Kim et al., 2010). Additionally, such enhanced hepatic P450 protein stability can also result in increased drug-elicited toxicity. Thus, the functional stabilization of hepatic Cyp2e1 rendered subtoxic concentrations of APAP, a popular over-the-counter analgesic and a Cyp2e1 substrate^2^, significantly more hepatotoxic to gp78^−/−^-hepatocytes than to corresponding WT.

However, such a functional P450 protein stabilization can also be beneficial. Thus, our finding of the enhanced bioactivation of the breast cancer chemotherapeutic prodrug tamoxifen in gp78^−/−^-relative to WT-hepatocytes, is particularly relevant and noteworthy. The E3 Ub-ligase gp78/AMFR was originally identified as the cell surface receptor for the prometastatic factor, autocrine motility factor (AMF; Nabi et al., 1990). Later, it was found to function as the E3 Ub-ligase in the ERAD of several substrates, including the tumor metastatic suppressor KAI1 (Fang et al., 2001; Tsai et al., 2007; Chen et al., 2012). Indeed, gp78 levels are highly upregulated within various human tumors (including lung, breast and colon) and their microenvironments, but not in adjacent normal tissue (Chiu et al., 2008; Fairbank et al., 2009). Importantly, this gp78-upregulation correlates with their invasiveness, the severity of their metastatic progression and poor prognosis (Chiu et al., 2008; Fairbank et al., 2009). Thus, African-American breast cancer patients despite their lower breast cancer incidence, exhibit markedly aggressive breast cancer forms and higher mortality rates than their European-American counterparts with lower mortality rates, albeit much higher breast cancer incidence (Chiu et al., 2008; Martin et al., 2009). Gene profiling via mRNA expression analyses revealed that gp78/AMFR is one of the two genes differentially elevated in the breast tumor epithelia and/or stroma of African-American patients versus those of their European-American cohorts. IB analyses documented a correspondingly increased gp78 protein content in these tissues (Martin et al., 2009). Similarly, a transgenic mouse mammary gland gp78-overexpression model also exhibited a hyperplastic phenotype, increased ductal branching and dense alveolar lobule formation that correlated with its downregulation of the KAI1 metastatic suppressor (Joshi et al., 2010). Thus, given our findings of enhanced Cyp3a-mediated tamoxifen bioactivation upon gp78-KO (Fig. 9), pharmacological attenuation of gp78 function may be therapeutically desirable as an adjuvant to concurrent breast cancer chemotherapy with prodrugs (such as tamoxifen) that are bioactivated by P450s. Plausible adjuvants include aspirin, salicylates and APAP that activate p38 MAPK-elicited gp78-degradation, resulting in CYP3A23 stabilization (Ohtsuki et al., 2019). However, although such a short-term therapeutic gp78-inhibition strategy warrants consideration, any long-term strategy may be contraindicated given that aged *gp78*-*null* mice reportedly develop NASH and human hepatocarcinoma (HHC) (Zhang et al., 2015).

Over 600 known mammalian E3 Ub-ligases exist that are functionally redundant in substrate-recognition and targeting (Metzeger et al., 2014), and capable of serving in both E3 and E4 (poly-Ub chain elongation of existing monoubiquitinated residues) complementary roles (Morito et al., 2008; Wang et al., 2013; Zhang et al., 2015; Menzies et al., 2018). Remarkably, despite these well-recognized E3-features, the genetic ablation of just a single hepatic E3, gp78/AMFR, is sufficient to stabilize its target substrates i.e. P450s, with significant pharmacological/toxic consequences.

Indeed, we recently documented that upon genetic CHIP-ablation in mice, hepatic Cyp2e1 was functionally stabilized irrespective of the presence of gp78 and possibly other redundant/complementary E3 Ub-ligases. This underscored the principal role of UbcH5a/CHIP/Hsp70 as the relevant E3-complex in Cyp2e1 proteolytic turnover (Kim et al., 2016a). This functional Cyp2e1 stabilization *in vivo* led to progressively increased hepatic lipid peroxidation and oxidative stress, JNK-activation, initial hepatic microvesicular steatosis that upon aging developed into macrovesicular steatosis, ushering a full-blown NASH syndrome over a one year-period (Kim et al., 2016a). These findings revealed that abrogation of CHIP-mediated Cyp2e1 ubiquitination not only had acute therapeutic/pharmacological consequences, but also severe, long-term pathophysiologic sequelae in mice.

Surprisingly, Cyp2e1-functional stabilization upon liver conditional gp78-KO although of a comparable magnitude and toxicologically significant, did not result in a similar hepatic microvesicular steatosis (D. Kwon & M. A. Correia, preliminary findings). In fact, fructose-induced hepatic triglyceride (TG) and free fatty acid content was significantly lower in gp78^−/−^ relative to gp78^+/+^-hepatocytes. This may stem from two plausible synergistic scenarios: (i) Concurrent stabilization of hepatic Insigs 1 and 2 upon gp78-KO may have suppressed SREB-elicited hepatic lipid biosynthesis as reported in a similar liver-conditional gp78-KO mouse model fed an obesity-inducing diet (Liu et al., 2012); and/or (ii) the concurrent stabilization upon gp78-KO of apolipoprotein B 100, an established gp78-substrate (Liang et al., 2003), critically involved in hepatic VLDL/LDL-packaging and export, may have accelerated extrahepatic TG-export, thereby minimizing the potential for hepatic steatosis. Our observations and those of others (Liu et al., 2012) reveal that upon their liver-conditional gp78-deletion, the mice are leaner and healthier, whereas global genetic gp78-deletion enhances their risk of hepatic steatosis, NASH and HCC, (Zhang et al., 2015).

While the gp78/AMFR upregulation in cancerous tissues is widely documented and unequivocal (Chiu et al., 2008; Martin et al., 2009), less clear is whether genetic gp78-polymorphisms normally exist in the general population, engendering correspondingly high and low (defective) gp78 functional phenotypes. The PubMed online human AMFR Gene/dbSNP Variant/Clin Variant databases list various gp78/AMFR allelic variants, and classify some as benign and others as pathologic. Arg/Arg mutations to Arg/Gly or Gly/Gly at “locus 125/145” are claimed to correlate with a higher expression of gp78 mRNA/protein, and correspondingly increased incidence of coronary atherosclerotic disease in a Chinese population (Wang et al., 2018b)^3^. The higher incidence of neuroglioblastoma multiformis in an Iranian population has also been associated with an AMFR genetic polymorphism (Eishi Oskouei et al., 2018). The validity of these claims remains however to be verified more extensively. Although, the gp78 SNPs listed in the PubMed Database are present along its entire 643 amino acid length, intriguingly, a few pertaining to basic (K, R) residues in the functional gp78 cytoplasmic domain, target the very same positively charged residues that we documented through chemical crosslinking and site-directed mutagenesis to be involved in its molecular recognition of CYP3A4 and CYP2E1 acidic phosphodegrons (Wang et al., 2015). Whether any of these listed SNPs truly contribute to genetic gp78-polymorphisms in human populations and/or correspondingly alter hepatic P450 content and function triggering DDIs, remains to be elucidated.

In summary, our findings document that the liver-specific genetic ablation of the E3 Ub-ligase gp78/AMFR can result through ERAD disruption in concurrent stabilization of several therapeutically relevant P450s with potentially serious, unwarranted therapeutic and/or toxic consequences. Such outcomes could be further aggravated in patients requiring multiple drug therapies for their specific conditions. Whereas for most drugs, their accelerated elimination and consequently shortened pharmacological response upon such P450-stabilization may be largely undesirable, this feature could also be ingeniously exploited as a therapeutic strategy in certain instances to enhance the metabolic activation of chemotherapeutic prodrugs with low therapeutic indices.

## Abbreviations used

APAP: Acetaminophen
ALT: alanine aminotransferase
AMF: autocrine motility factor
AMFR: AMF-receptor
ALD: autophagy-lysosomal degradation
β-NF: β-naphthoflavone
CHIP: Carboxy-terminus of Hsc70-interacting protein
CHX: cycloheximide
P450s; CYPs: cytochromes P450
DEX: dexamethasone
DDIs: drug-drug interactions
ER: endoplasmic reticulum
ERAD: ER-associated degradation
IB: immunoblotting
INH: isoniazid
KO: knockout
3-MA: 3-methyladenine
NASH: nonalcoholic steatohepatitis
PB: phenobarbital
shRNAi: shRNA interference
TG: triglyceride
Ub: ubiquitin
UPD: Ub-dependent proteasomal degradation
WT: wild type

## Acknowledgments

We thank Mr. Chris Her for liver cell isolation at the UCSF Liver Center Core on Cell & Tissue Biology, supported by NIDDK Center Grant DK26743. We thank Dr. Alexandra Ioanoviciu for the training in the use of LC-MS system used in the tamoxifen metabolite analyses. We greatly appreciate the contributions of Lisa Yu and Trisha Mao for LC-MS/MS analyses of nicotine metabolites in cell culture samples, and funding from NIDA P30 DA012393 (to N. Benowitz, UCSF) for supporting the laboratory resources used for these analyses.

## Footnotes

a) This work was supported by NIH Grants GM44037 and DK26506 to M. A. C.

b) This was presented in poster form and in one of the ASPET “Data Blitzes” at the 2018 Experimental Biology, Abstract C148.

c) Reprint requests to M. A. Correia.

## Footnotes

1 This is also true of the other P450-dependent cotinine metabolites, whose relative increases were maximally detected at 24 h. By contrast, the FMO3-metabolite nicotine-N-oxide was decreased, rather than increased in gp78^−/−^-hepatocytes, thereby revealing that FMO3 was, if at all, decreased rather than increased upon gp78-KO.

2 Although, without concurrent Cyp3a-induction in cultured INH-pretreated gp78^−/−^-hepatocytes Cyp3a levels are low, gp78-ablation would substantially synergize such toxicity *in vivo* due to their functional stabilization.

3 The authors indicate that the Arg/Arg residues at “locus” 145 or 125 were affected by the polymorphism. However, the corresponding residues in the human gp78/AMFR primary sequence deposited in PubMed are Val145 and Phe125.

